# Comparison of Four Density Gradient Centrifugation Protocols for Purification of *Toxoplasma gondii* PRU Brain Cysts

**DOI:** 10.64898/2026.06.21.733578

**Authors:** Tingting Yi, Wenyu Ma, Xuefei Liang, Shuaijing Niu, Jian An, Huanrong Li, Qiuming Li, Deqi Yin

**Affiliations:** Coccidia Laboratory, College of Animal Science and Technology/College of Veterinary Medicine, Beijing University of Agriculture, Beijing 102206, China

**Keywords:** *Toxoplasma gondii*, brain cyst, density gradient centrifugation, isolation and purification, viability

## Abstract

*Toxoplasma gondii* is an obligate intracellular zoonotic protozoan that establishes persistent brain cysts in infected hosts, causing chronic infection and neuropathological damage. Efficient isolation and purification of brain cysts are essential for studying its biological characteristics and developing effective control strategies. The present study aimed to establish a reliable method for purifying brain cysts of the *T. gondii* PRU strain with high yield, viability, and practicality. Brain cysts harvested from experimentally infected ICR mice were purified using four different density gradient centrifugation methods: lymphocyte separation medium (LSM), Percoll, cesium chloride (CsCl), and sucrose. Purification efficiency was systematically evaluated, and the infection model was validated via brain histopathology. Cyst counts and purification yields were quantified, while cyst and bradyzoite viability were assessed using FDA/PI staining, trypan blue exclusion, and in vivo infectivity assays. Infected mice displayed the most severe clinical signs at 15 days post-infection (dpi), accompanied by significantly reduced body weight compared with uninfected controls (*P <* 0.05) and prominent perivascular inflammatory infiltration in the brain. Among the four purification methods, Percoll, CsCl, and sucrose gradients yielded significantly higher cyst numbers than LSM (*P <* 0.001), with no significant differences observed among the three gradient-based methods. Cysts purified by Percoll and LSM retained high viability and remained fully infectious in mice. Sucrose-purified cysts exhibited decreased viability, but their bradyzoites remained infective. In contrast, cysts and bradyzoites purified by CsCl were completely non-viable. These results clarify the advantages and limitations of each protocol and provide an optimized technical reference for *T. gondii* brain cyst research, supporting the development of toxoplasmosis prevention and control measures.

## Introduction

*Toxoplasma gondii* is an obligate intracellular parasite with a worldwide distribution, able to infect nucleated cells of almost all warm-blooded vertebrates (Ji et al. 2026). Epidemiological statistics indicate that approximately one-third of the global human population has been infected with this pathogen. In immunocompetent hosts, infection predominantly progresses to a chronic state, characterized by persistent encystment of the parasite within neurons to establish immune-privileged niches (Schlüter & Barragan 2019). Upon deterioration of host immune competence, these tissue cysts reactivate and rupture, potentially triggering fatal toxoplasmic encephalitis (Maus et al. 2024; Wang et al. 2026). To date, no approved chemotherapeutics or efficacious vaccines are available for the treatment of chronic toxoplasmosis. As the predominant parasitic form during chronic *T. gondii* infection, brain cysts serve as core experimental substrates for drug screening, vaccine development, and mechanistic investigations into parasite pathogenesis. Accordingly, the efficient isolation and purification of intact, viable brain cysts constitute an essential prerequisite for in vitro research on chronic toxoplasmosis.

In the 1960s, Nakabayashi and Motomura (1968) first applied gum arabic multi-layer centrifugation and sucrose density gradient centrifugation to isolate *T. gondii* cysts from mouse brain, establishing early protocols for cyst purification. However, gum arabic-based methods suffer from low throughput and lengthy processing, limiting their large-scale application(Nakabayashi & Motomura 1968). Subsequently, Cornelissen et al. (1981) developed continuous Percoll density gradient centrifugation, which improves separation efficiency and purity while yielding high cyst recovery and maintaining cyst viability; this method has become the mainstream isolation approach in this field (Watts et al., 2017). (Watts et al. 2017). Lymphocyte separation medium (Ficoll-Paque) was later validated for cyst isolation due to its technical simplicity (Li et al. 2024). Cesium chloride (CsCl) gradient centrifugation proved effective for purifying *T. gondii* oocysts and tachyzoites, as well as *Cryptosporidium* oocysts (Staggs et al. 2009; Chen and Huang. 2006). Nonetheless, a systematic evaluation of CsCl for brain cyst purification is still lacking, and its applicability remains poorly defined. Previous purification protocols were validated under disparate experimental setups with inconsistent parameters, and few studies have conducted standardized quantitative assessments of cyst recovery, parasite viability and assay reproducibility. Herein, we systematically compared four density gradient centrifugation approaches, namely lymphocyte separation medium, Percoll, sucrose and cesium chloride (CsCl), under fully identical experimental conditions. We comprehensively assessed their purification performance with respect to cyst yield, sample purity, bradyzoite viability, experimental cost and operational feasibility, and facilitate mechanistic studies and anti-*toxoplasma* drug development against chronic toxoplasmosis.

## Results

### Clinical Observations and Body Weight Monitoring in PRU-Infected Mice

The workflow of brain homogenate preparation is shown in Fig. 1A. Infected mice exhibited ruffled fur from 7 days post-infection (dpi), with clinical manifestations peaking at approximately 15 dpi, including piloerection, decreased locomotor activity, incomplete eyelid closure, and hunched posture. These symptoms gradually alleviated and resolved by 30 dpi (Fig. 1B). Clinical scores (higher values indicate worse health status) are presented in Fig. 1C. A 40-day body weight monitoring showed that the infected group displayed a significant weight reduction starting at 15 dpi relative to the control group (Fig. 1D).

**Fig. 1.**
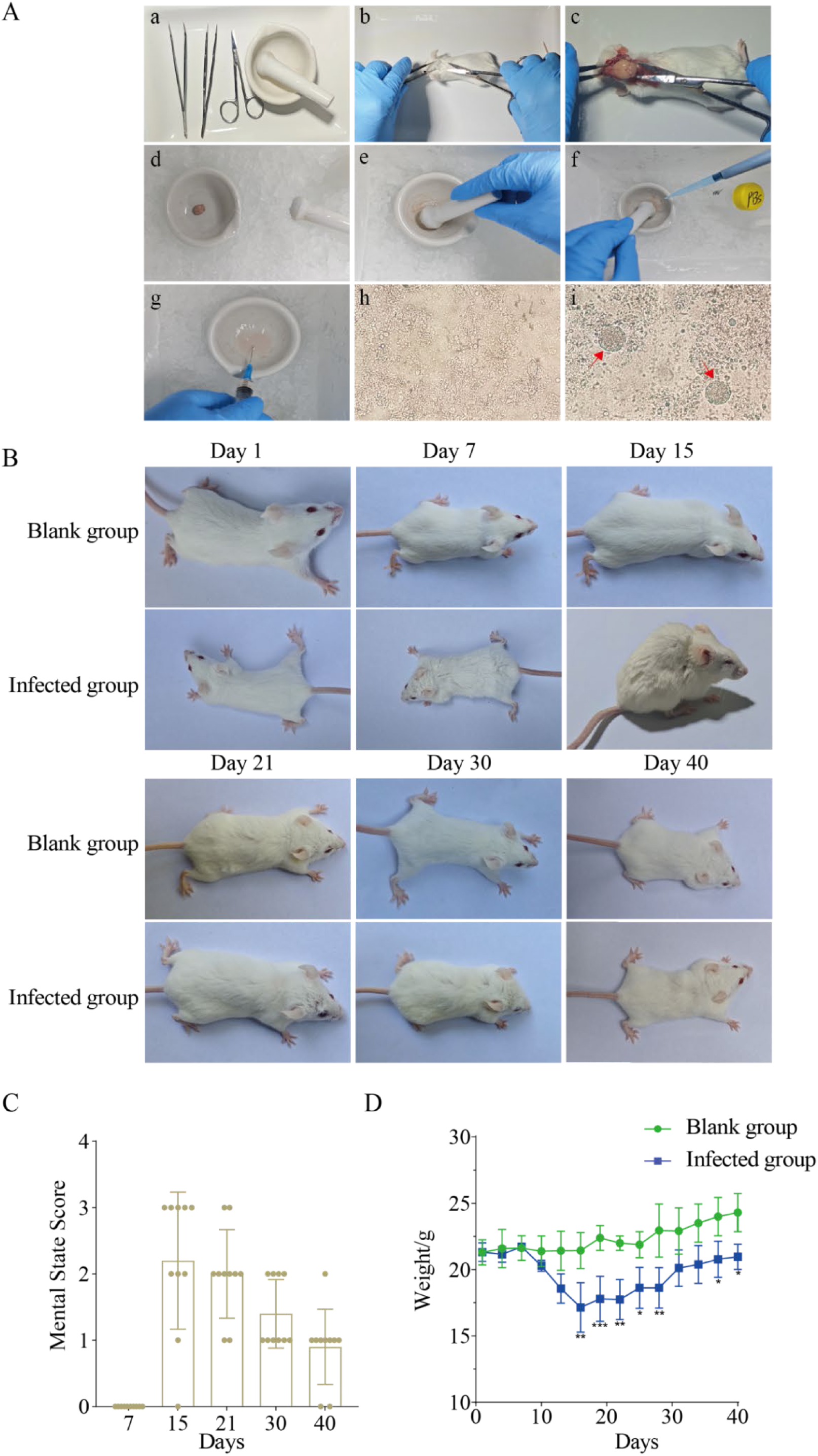
Establishment of the *T. gondii* PRU chronic infection model and brain cyst observation A: Brain homogenate preparation and microscopic examination. (a) Dissecting tools and mortar; (b) Incision of the scalp between the ears; (c) Removal of the parietal bone and isolation of whole brain; (d) Rinsing and transfer of brain tissue to a pre-chilled mortar; (e) Gentle grinding into a paste; (f) Addition of 1% Tween-80-PBS; (g) Repeated aspiration with a 23G syringe; (h) Microscopic view of control brain homogenate; (i) Microscopic view of infected brain homogenate. B: Clinical appearance of infected mice. C: Clinical scoring of infected mice. D: Body weight changes of mice.

### Histopathological Analysis of Mouse Brain Tissue

Histopathological examination revealed no pathological lesions in uninfected mouse brains. In contrast, infected brains exhibited typical *Toxoplasma gondii*-associated pathological features. Hematoxylin and eosin (HE) staining identified intact *T. gondii* cysts filled with bradyzoites within the brain parenchyma. Prominent neuronophagia was observed, with microglia infiltrating and phagocytosing necrotic neurons and their processes. Characteristic perivascular cuffing, defined by perivascular accumulation of lymphocytes and glial cells, was also evident (Fig. 2).

**Fig. 2.**
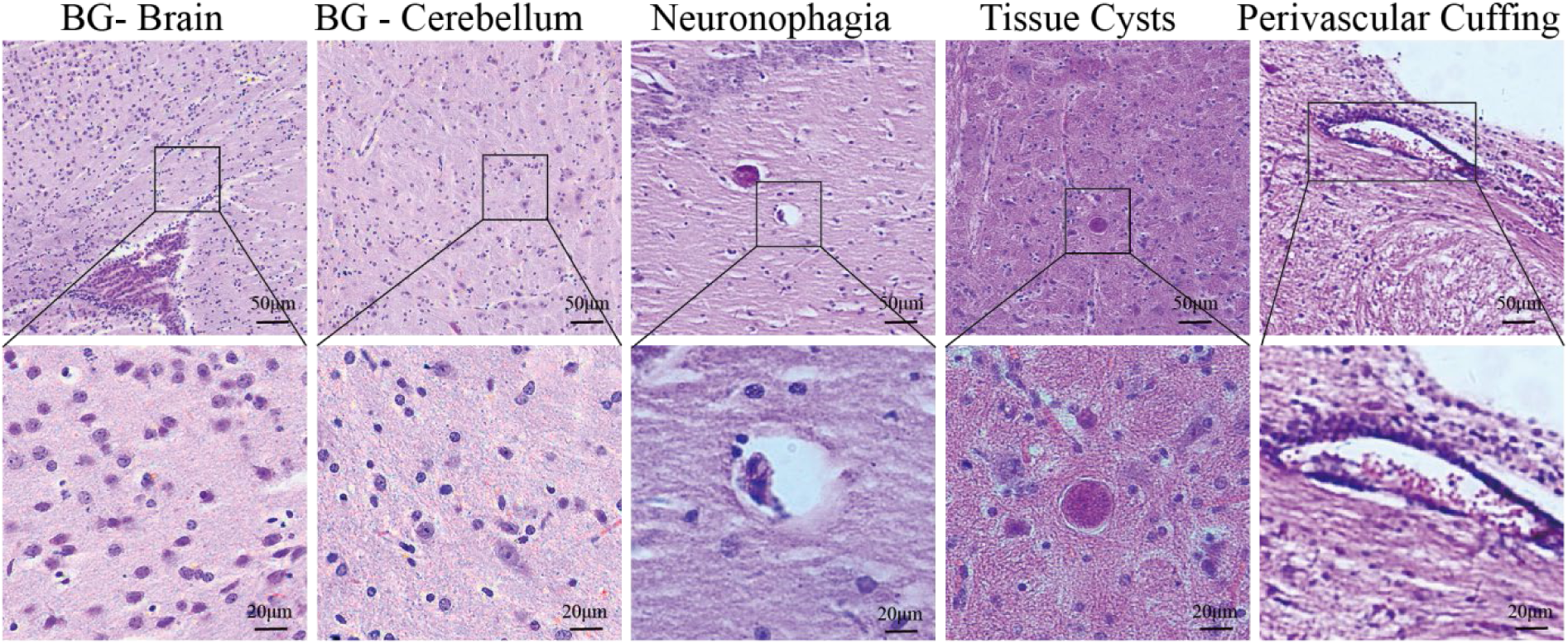
Histopathological observation of mouse brain tissue (HE staining)

### Purity Comparison of Purified Brain Cysts

All four density gradient centrifugation methods produced distinct, clear separation interfaces (Fig. 3A, 3E, 3I, 3M). Microscopic analysis revealed marked differences in residual debris among the purified preparations. The lymphocyte separation medium (LSM) group contained an average of 15 tissue fragments per field (Fig. 3C, 3D); the Percoll group had 7 fragments per field (Fig. 3G, 3H); the sucrose group averaged 5 fragments per field (Fig. 3K, 3L); and the cesium chloride (CsCl) group showed the least debris, with the highest solution clarity and optimal purification efficiency (Fig. 3O, 3P).

**Fig. 3.**
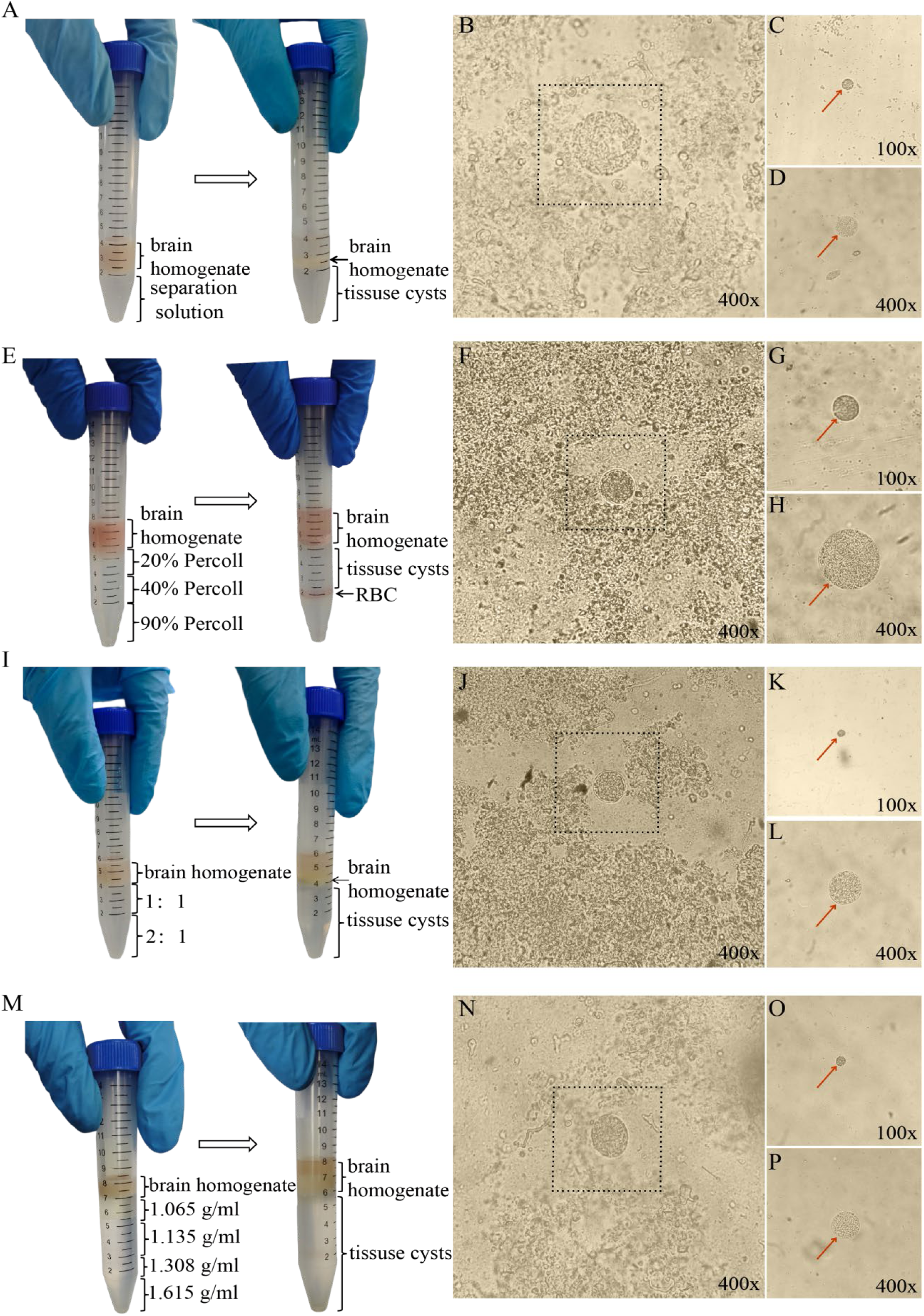
Gradient setup and microscopic examination of brain cysts before and after purification. A, E, I, M: Schematic diagrams showing density gradient stratification of Ficoll-Paque lymphocyte separation medium, Percoll, sucrose, and cesium chloride (CsCl), respectively. Brain homogenate was loaded onto the top of each gradient. B, F, J, N: Micrographs of *T. gondii* cysts present in unpurified brain homogenate from each group. C, D, G, H, K, L, O, P: Representative images of purified *T. gondii* brain cysts isolated using the four aforementioned methods; panels C, G, K, O were captured at 100× magnification, whereas D, H, L, P were captured at 400× magnification.

### Cyst Yield and Recovery of Four Purification Methods

Purification yields differed significantly among the four methods. The mean cyst counts were: lymphocyte separation medium (LSM), (0.92 ± 0.001) × 10³; Percoll density gradient centrifugation, (1.01 ± 0.003) × 10³; sucrose density gradient centrifugation, (1.07 ± 0.002) × 10³; and cesium chloride (CsCl) density gradient centrifugation, (1.12 ± 0.003) × 10³. The corresponding recovery rates were 61.5%, 70.0%, 71.6%, and 72.0%, respectively. Statistical analysis revealed that the yields of Percoll, sucrose, and CsCl methods were significantly higher than that of LSM (*P <* 0.001), with no significant differences among the three gradient-based methods (Table 1).

**Table 1.**
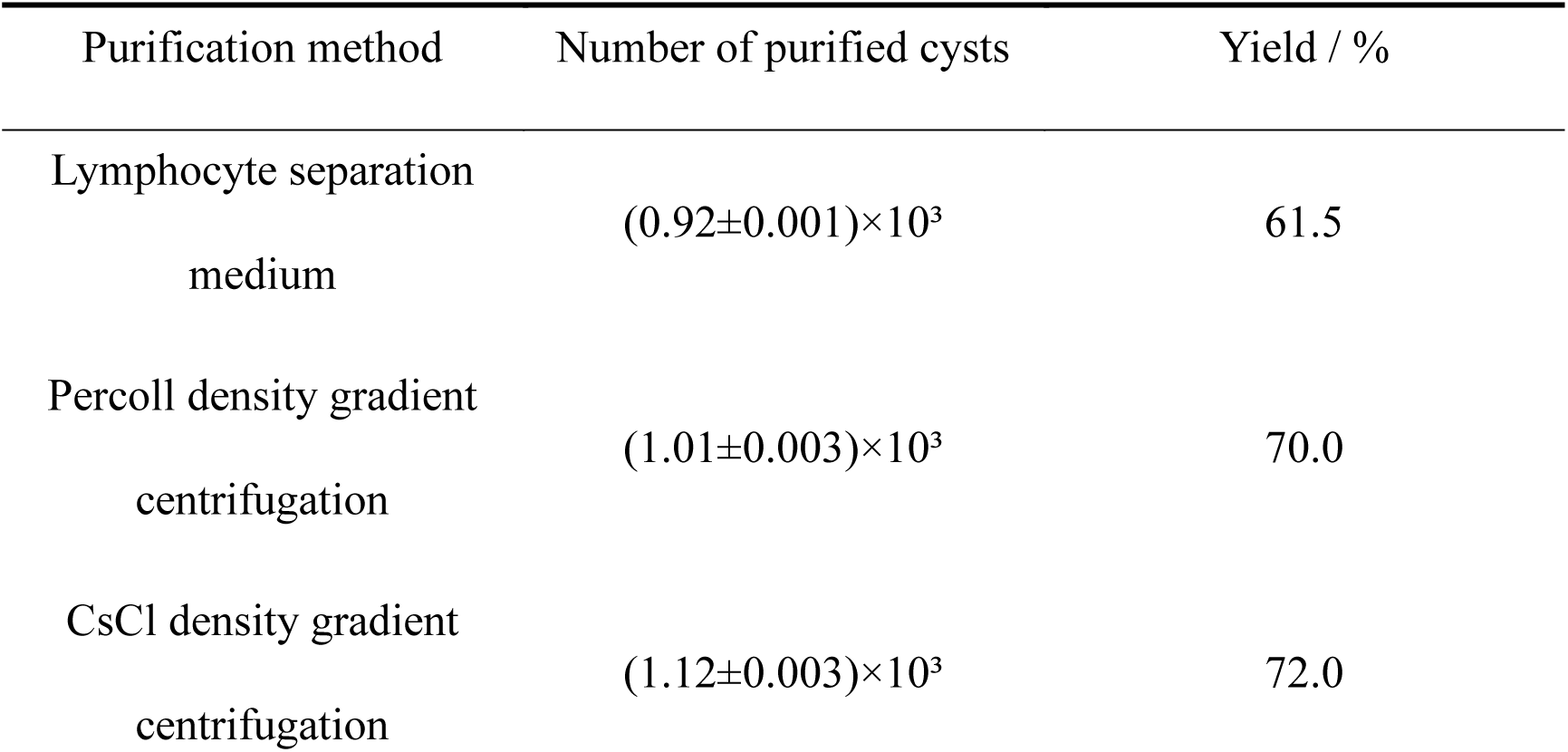
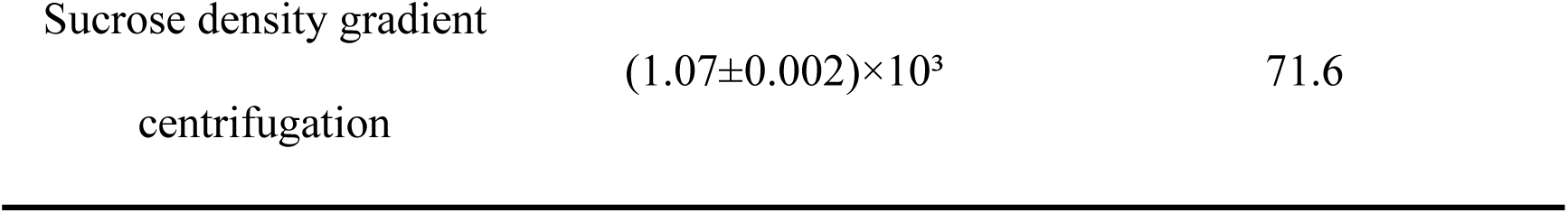
Comparison of Purification Yields of Four Methods for *Toxoplasma gondii* Brain Cysts.

### Viability Assessment of Purified Cysts by Mouse Infection and FDA/PI Staining

Following in vivo infection, brain cysts were recovered from mice inoculated with cysts purified by Percoll density gradient centrifugation or lymphocyte separation medium (LSM), but not from those receiving sucrose- or cesium chloride (CsCl)-purified cysts. Fluorescein diacetate/propidium iodide (FDA/PI) staining confirmed that Percoll- and LSM-purified cysts exhibited green fluorescence, indicating high viability. In contrast, most bradyzoites within sucrose- and CsCl-purified cysts showed red fluorescence, reflecting markedly reduced or complete loss of viability (Fig. 4).

**Fig. 4.**
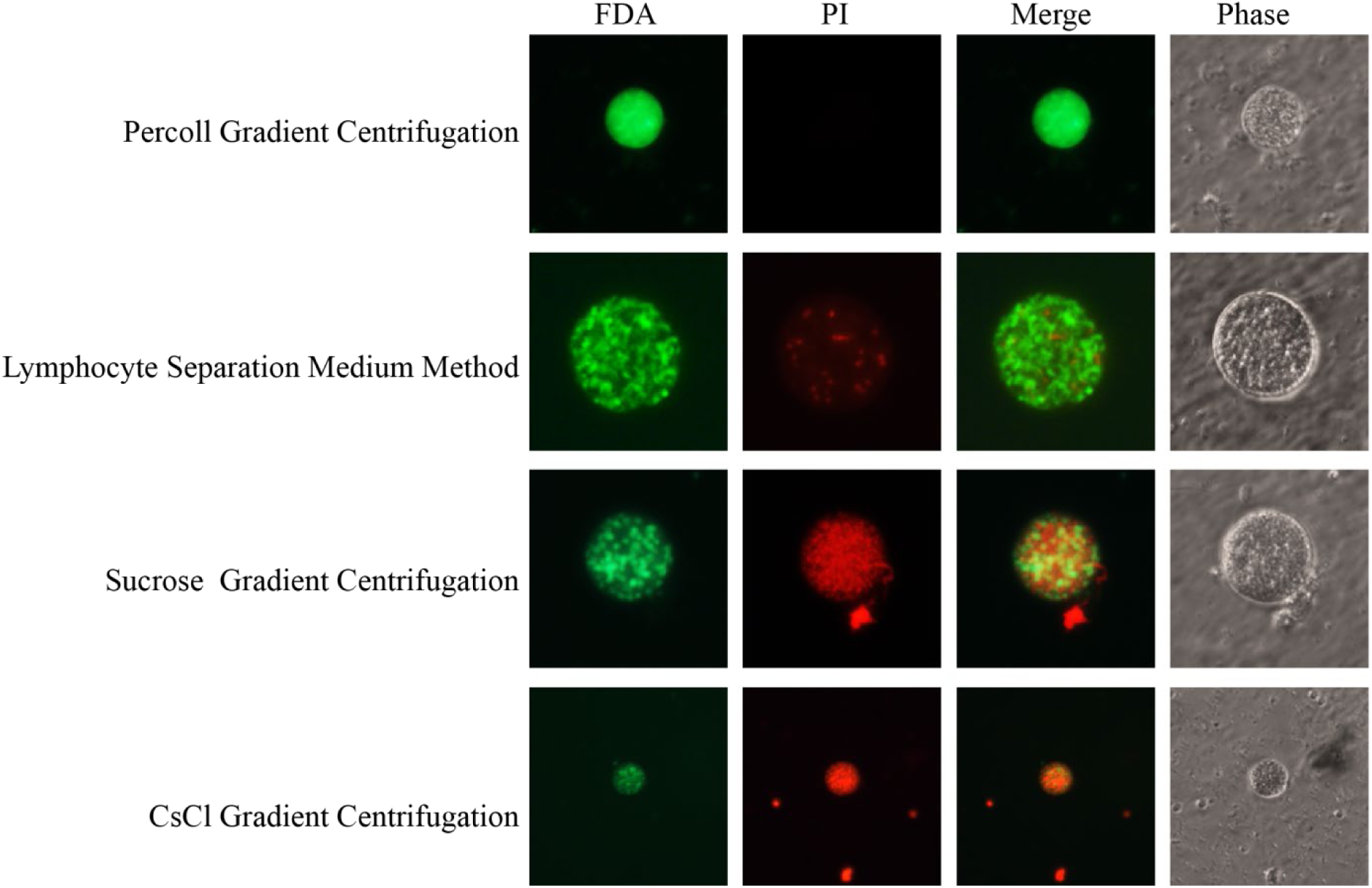
Viability staining of brain cysts purified by four methods (200×)

### Assessment of Bradyzoite Viability by Trypan Blue Staining

Trypan blue exclusion assays revealed that purification method significantly affected bradyzoite viability. Percoll density gradient centrifugation yielded the highest viability (98%), followed by lymphocyte separation medium (95%), sucrose density gradient centrifugation (75%), and cesium chloride density gradient centrifugation (55%) (Fig. 5).

**Fig. 5.**
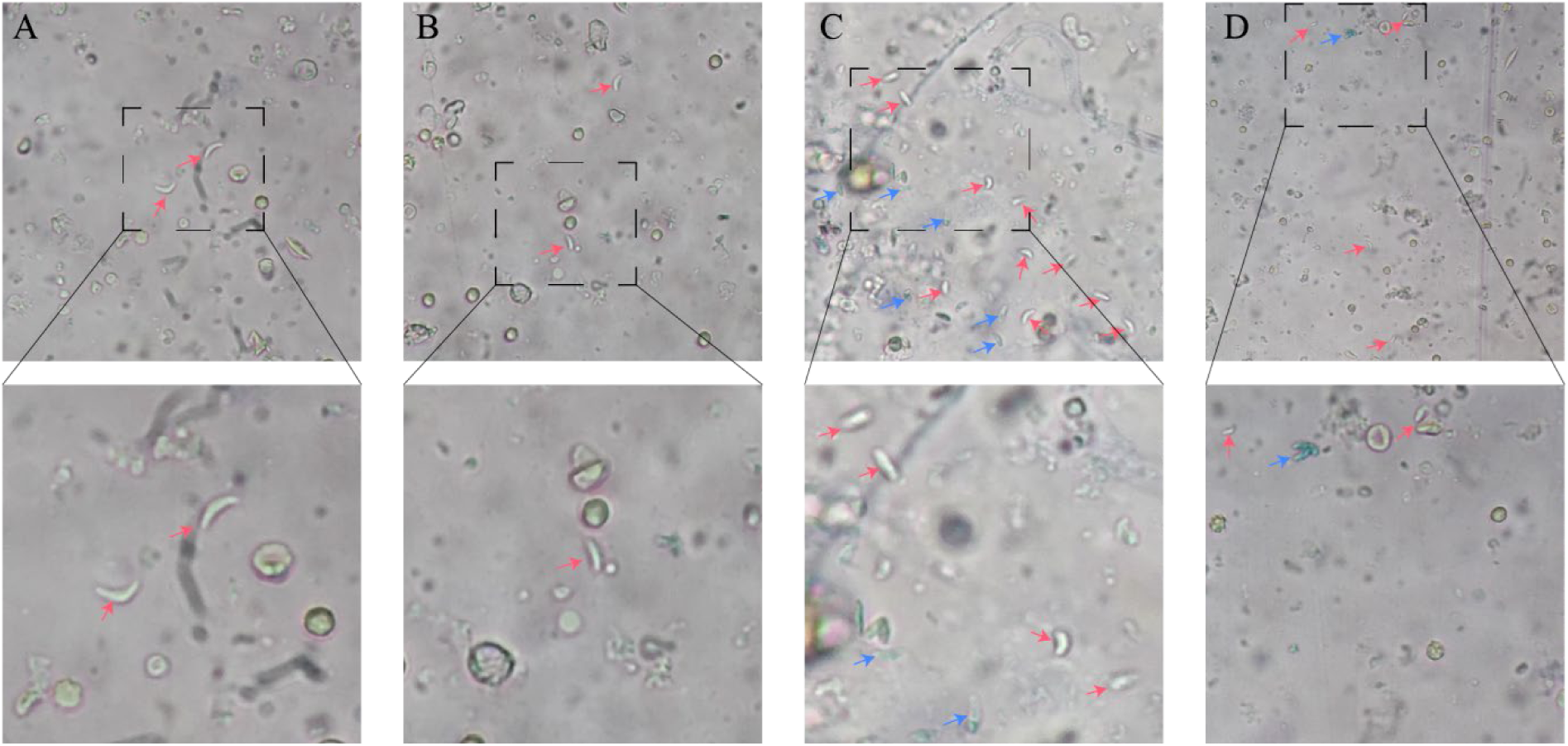
Trypan blue staining of bradyzoites in purified brain cysts (200×) A: Percoll density gradient centrifugation; B: Lymphocyte separation medium (LSM); C: Sucrose density gradient centrifugation; D: Cesium chloride (CsCl) density gradient centrifugation. Red arrows mark viable *T. gondii*, and blue arrows mark non-viable parasites.

### Infectivity of Isolated Bradyzoites in Mice

Mice were intraperitoneally inoculated with 500 bradyzoites each. At 30 days post-infection (dpi), five mice from each group were randomly euthanized, and their brain tissues were examined microscopically. Typical *T. gondii* cysts were detected in all groups except the cesium chloride (CsCl) group, where no cysts were found (Fig. 6). Quantitative analysis showed the average cyst loads per mouse were 800 in the Percoll group, 733 in the lymphocyte separation medium (LSM) group, and 600 in the sucrose group.

**Fig. 6.**
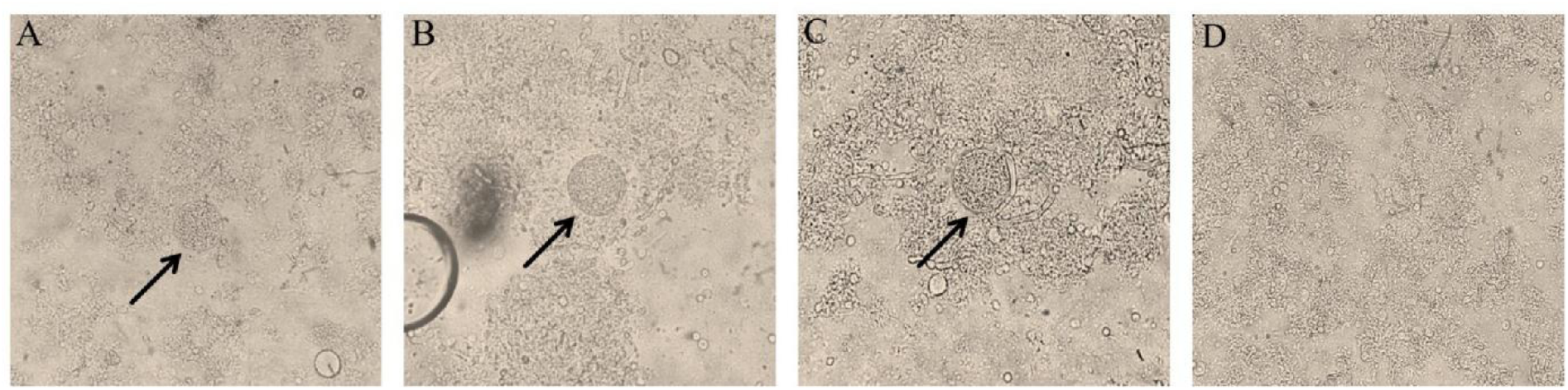
Microscopic view and quantification of brain cysts after bradyzoite infection (400×) A: Percoll density gradient centrifugation; B: Lymphocyte separation medium (LSM); C: Sucrose density gradient centrifugation; D: Cesium chloride (CsCl) density gradient centrifugation.

### Cost Comparison of Four Purification Methods

We further compared the overall costs of the four protocols for purifying *T. gondii* brain cysts. In terms of reagent expenses, cesium chloride (CsCl), lymphocyte separation medium (LSM) and Percoll incurred higher per-test costs, while sucrose was the most economical option. For equipment requirements, the CsCl method needed a refrigerated low-speed centrifuge, leading to extra expenditure, whereas the other three approaches only used a standard low-speed centrifuge.

In terms of operation, the CsCl protocol was the most labor-intensive, as it required preparation of four distinct gradients and corresponding buffers. Percoll and sucrose methods presented moderate technical difficulty, with only dilution needed to form discontinuous gradients. By contrast, the LSM method was the easiest to operate, since commercial reagents could be used directly for purification. In terms of time consumption, the CsCl method took the longest processing time. Percoll and sucrose groups had comparable time costs, and the LSM method was the most time-efficient. Detailed data are summarized in Table 2.

**Table 2.**
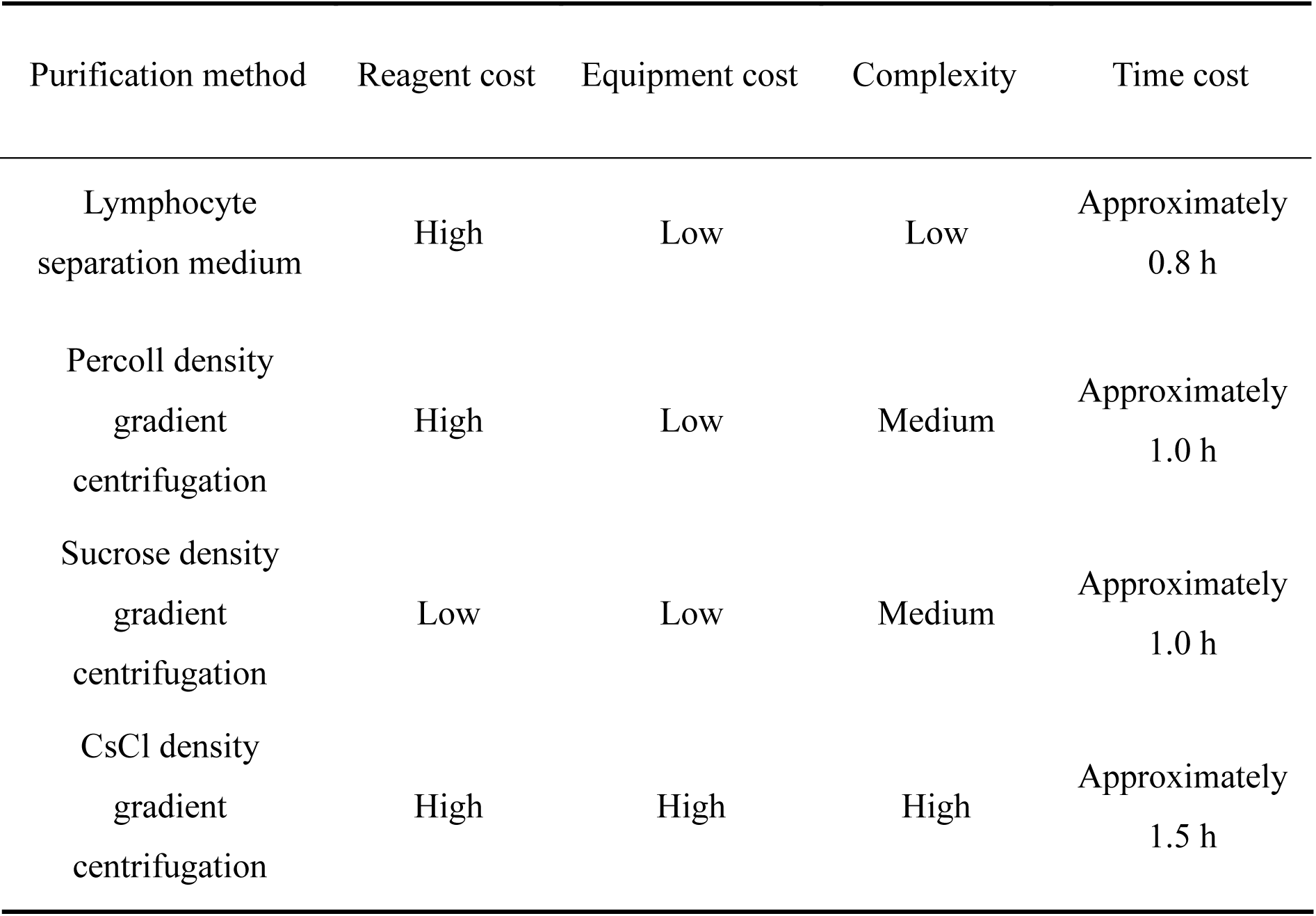
Cost and operational comparison of four purification methods.

| Purification method | Reagent cost | Equipment cost | Complexity | Time cost |
| --- | --- | --- | --- | --- |
| Lymphocyte<br>separation medium | High | Low | Low | Approximately<br>0.8 h |
| Percoll density<br>gradient<br>centrifugation | High | Low | Medium | Approximately<br>1.0 h |
| Sucrose density<br>gradient<br>centrifugation | Low | Low | Medium | Approximately<br>1.0 h |
| CsCl density<br>gradient<br>centrifugation | High | High | High | Approximately<br>1.5 h |

## Discussion

In the present study, four mainstream density gradient centrifugation protocols involving LSM, Percoll, sucrose, and CsCl were systematically compared under unified and standardized experimental conditions for purifying *T. gondii* PRU strain brain cysts. Comprehensive evaluation covering purification yield, sample purity, parasite viability, experimental cost, and operational feasibility confirmed that Percoll density gradient centrifugation is the most balanced and reliable purification method. Consistent with previous classic studies (Cornelissen et al. 1981; Watts et al. 2017) , Percoll-based purification enables efficient separation of intact brain cysts from mouse brain homogenates while retaining high cyst recovery and bradyzoite viability. Benefiting from its isotonic separation environment, Percoll avoids severe osmotic shock and chemical damage to parasites, showing unique superiority in the preparation of viable *T. gondii* cysts for biological functional experiments. This method provides a standardized and stable technical scheme for subsequent researches on the biological characteristics, pathogenic mechanisms, and drug screening of *T. gondii* brain cysts, and strongly supports the technical development of toxoplasmosis prevention and control. FDA/PI double fluorescence staining intuitively reflected the differential damage of different purification media to cyst and bradyzoite activity. Cysts purified by Percoll and LSM presented only green fluorescence, indicating intact bradyzoite membrane structure and normal intracellular esterase activity, which are core indicators of high biological viability (Wang et al. 2025). The mild biochemical properties of these two media may be the key to maintaining parasite viability. Percoll operates under strict isotonic conditions, which fundamentally eliminates osmotic stress-induced structural damage to parasites. Similarly, commercial LSM can effectively buffer osmotic pressure changes and reduce non-specific chemical toxicity, thus completely preserving the structural integrity and physiological function of bradyzoites (Pertoft 2000). In contrast, sucrose and CsCl gradient purification groups exhibited mixed red and green fluorescence, demonstrating partial membrane permeabilization while residual metabolic activity remained. Both hypertonic sucrose solution and CsCl solution possess cell permeability. Notably, cesium ions have been proven to exert significant dose-dependent cytotoxic effects on eukaryotic cells by interfering with intracellular metabolic pathways and ion balance (Kobayashi et al. 2017). Therefore, we speculate that the high osmotic pressure formed by sucrose and CsCl gradients increases the permeability of bradyzoite cell membranes, allowing external solutes to invade the parasite, causing partial structural damage and metabolic imbalance, which ultimately leads to the mixed staining phenotype.

The intact membrane of bradyzoites determines their resistance to digestion by gastric acid and pepsin in the host gastrointestinal tract, which is critical for the oral infectivity and transmission of *T. gondii* tissue cysts (Dubey 1998). In this study, oral infection verification showed that sucrose-purified cysts completely lost oral infectivity, while bradyzoites isolated from sucrose-treated cysts still successfully established infection via intraperitoneal inoculation. This finding confirms that sucrose purification causes only reversible membrane injury to bradyzoites. The hypertonic sucrose environment weakens the parasite’s tolerance to gastrointestinal digestive enzymes, but does not cause irreversible cell death, and the intraperitoneal physiological microenvironment can repair mild membrane damage and restore parasite infectivity. Differently, CsCl-purified cysts completely lost infectivity through both oral and intraperitoneal inoculation routes. Combined with trypan blue staining results that nearly half of the bradyzoites were inactivated, it is verified that CsCl treatment induces irreversible cytotoxic damage to *T. gondii* bradyzoites. Importantly, our study further proves that single cytochemical staining methods including FDA/PI and trypan blue staining can only reflect the instantaneous membrane integrity and basic metabolic activity of parasites, which cannot fully represent the actual in vivo infectivity of cysts. Excessive reliance on in vitro staining results will significantly overestimate the survival rate of purified cysts. Therefore, in vivo animal infectivity assay is an indispensable gold standard for accurately evaluating the biological activity of *T. gondii* cysts and bradyzoites, which should be supplemented in all parasite viability evaluation experiments.

In conclusion, Percoll and LSM have minimal adverse impacts on the viability of cysts and bradyzoites, making them ideal for experiments that require high biological activity of parasites. Although sucrose density gradient centrifugation induces reversible membrane damage in bradyzoites, the parasites remain infectious via intraperitoneal inoculation. Given its low cost, this method is suitable for preparing highly purified cysts when oral infection is not required. By contrast, CsCl treatment causes irreversible damage and complete loss of infectivity, so it is not applicable for experiments using live cysts. Overall, each purification method has distinct applicable scenarios. Percoll density gradient centrifugation is the first choice when both high viability and high yield are required. LSM is preferable for simple and rapid purification of small-volume samples. Sucrose density gradient centrifugation is suitable for low-cost large-scale purification in studies without oral infection procedures. CsCl density gradient centrifugation can be adopted exclusively for isolating highly purified dead cysts for structural and compositional analyses.

In summary, the four density gradient centrifugation methods have distinct advantages, limitations and applicable scenarios for *T. gondii* brain cyst purification. Percoll and LSM purification have negligible adverse effects on cyst and bradyzoite activity, making them the preferred methods for preparing high-viability parasite samples for functional experiments. Sucrose density gradient centrifugation, with prominent cost advantages and simple operation, can obtain high-purity cysts. Although it causes reversible membrane damage to bradyzoites and impairs oral infectivity, the residual infectivity via intraperitoneal injection is retained, which is suitable for low-cost and large-scale sample preparation for non-oral infection experiments. As previously reported, CsCl gradient centrifugation can achieve extremely high purification purity for *T. gondii* parasitic samples, but its strong cytotoxicity completely destroys parasite activity and infectivity (Staggs et al. 2009). Thus, this method is only applicable for the isolation and preparation of dead cysts, and can be used for subsequent parasite ultrastructural observation and component analysis experiments. In practical experimental applications, researchers can select the optimal purification scheme according to experimental purposes, sample volume, cost budget and activity requirements. Among all methods, Percoll density gradient centrifugation comprehensively balances yield, purity, viability and operability, and is the most universal and reliable technical scheme for routine purification of *T. gondii* brain cysts.

This study was performed using only the *T. gondii* PRU strain and ICR mice, which imposes certain limitations on our conclusions. Further validations involving other parasite strains, cysts derived from different host species, and long-term preservation of purified cysts were not conducted. Future work should include more parasite strains and host models, optimize protocols for the preservation and reactivation of purified cysts, and establish a comprehensive technical system for the isolation, purification and viability maintenance of *T. gondii* brain cysts. These efforts will provide reliable experimental materials for exploring the pathogenesis and immune evasion mechanisms of chronic toxoplasmosis, as well as for developing novel anti-*toxoplasma* drugs and vaccines.

## Conclusions

In summary, this study comprehensively assessed four protocols for the isolation and purification of *T. gondii* brain cysts. Taking yield, purity, viability, cost and practicality into consideration, Percoll density gradient centrifugation stands out as the optimal method. Our results provide a standardized methodological reference for relevant experiments, and lay a foundation for basic research on toxoplasmosis as well as the development of prevention and control technologies.

## Materials and Methods

### Parasite Strain, Animals, Reagents and Equipment

The *Toxoplasma gondii* PRU strain was routinely maintained and passaged in the Coccidia Laboratory, Beijing University of Agriculture.

#### Reagents

Percoll (P8370, Solarbio, China); lymphocyte separation medium (P8650, Solarbio, China); sucrose (S818048, Macklin, China); cesium chloride (C80679, Macklin, China); fluorescein diacetate (FDA, H973007, Dibo, China); propidium iodide (PI, ST1569, Beyotime, China).

#### Equipment

High-speed refrigerated centrifuge; inverted fluorescence microscope (Eppendorf, Germany); bench low-speed centrifuge TD6 (Huxi, China); light microscope (Nikon, Japan).

#### Animals

Eight-week-old specific pathogen-free (SPF) female ICR mice were purchased from Sibafu (Beijing) Biotechnology Co., Ltd. All mice were reared under standard laboratory conditions with ad libitum access to food and water. All animal experiments were approved by the Animal Ethics Committee of Beijing University of Agriculture (Approval No. BUA2023118). Mice were euthanized via cervical dislocation under deep isoflurane anesthesia upon experiment termination or when reaching humane endpoints.

### Experimental Design and Infection

Following a one-week acclimation period, 40 female ICR mice were randomly divided into the control group and infection group, with 20 mice per group. Mice in the control group were administered phosphate-buffered saline (PBS) via oral gavage, while each mouse in the infection group received an oral inoculation of 10 *T. gondii* cysts.

### Clinical Monitoring and Body Weight Assessment

Body weight was measured every two days throughout the experiment. Daily clinical evaluation was conducted according to fur appearance, locomotor activity, eyelid status and body posture. Photographic records were collected for phenotypic observation. The entire monitoring period lasted for 40 days.

### Microscopic Examination of Brain Cysts

Mice were euthanized at 40 days post-infection (dpi). Whole brain tissues were isolated, rinsed and homogenized in 2 mL ice-cold PBS supplemented with 1% Tween-80. The homogenate was repeatedly passed through a 23-gauge syringe needle 3 to 5 times. A 10 μL homogenate sample was observed under a light microscope at 10× magnification for cyst detection and enumeration.

### Brain Histopathological Analysis

Dissected brain tissues were fixed in 4% paraformaldehyde for 24 h. After gradient dehydration, tissue clearing and paraffin embedding, serial sections were prepared. The sections were deparaffinized, stained with hematoxylin and eosin (HE), dehydrated, cleared and mounted for microscopic observation.

### Preparation of Sucrose Density Gradient

The sucrose stock solution was prepared by dissolving 15.62 g sucrose in 10 mL double-distilled water (ddH₂O). Working solutions were subsequently obtained by diluting the stock solution with 1× PBS at volume ratios of 2:1 and 1:1.

### Preparation of Cesium Chloride (CsCl) Density Gradient

CsCl gradient solutions were formulated using Tris·HCl buffer. Briefly, 0.121 g Tris, 0.8 g NaCl and 0.037 g EDTA were dissolved in 90 mL ddH₂O. The pH was adjusted to 7.4 with concentrated hydrochloric acid, and the final volume was brought to 100 mL with ddH₂O. Detailed preparation protocols for CsCl gradients are listed in Table 3.

**Table 3.** Preparation of CsCl solutions with different gradients.

| Density | CsCl / g | Tris·HCl buffer / mL |
| --- | --- | --- |
| 1.065 g/mL | 0.13 | 2 |
| 1.135 g/mL | 0.27 | 2 |
| 1.308 g/mL | 0.615 | 2 |
| 1.615 g/mL | 1.23 | 2 |

### Percoll Gradient Preparation

Percoll working solutions were prepared with 10×PBS and ddH₂O as detailed in Table 4.

**Table 4.** Preparation of Percoll density gradients.

| Concentration | 10× PBS / mL | Percoll stock / mL | ddH<sub>2</sub>O / mL |
| --- | --- | --- | --- |
| 90% | 1 | 9 | 0 |
| 40% | 1 | 4 | 5 |
| 20% | 1 | 2 | 7 |

### Purification Using Lymphocyte Separation Medium

A total of 2 mL of brain homogenate was carefully layered onto 2 mL of lymphocyte separation medium (LSM) in a 15 mL centrifuge tube, followed by centrifugation at 1,280 g for 30 min at 4 °C. The upper supernatant was discarded, and the resulting pellet was washed with 10 mL of 1× PBS. After centrifugation at 1,580 g for 10 min, the pellet was resuspended in approximately 1 mL of PBS for cyst counting.

### Purification Using Sucrose Density Gradient Centrifugation

A discontinuous sucrose density gradient was constructed by sequentially layering 2 mL of 1:1 diluted sucrose solution and 2 mL of 2:1 diluted sucrose solution. Subsequently, 2 mL of brain homogenate was gently overlaid on the top of the gradient. After centrifugation at1280 g for 25 min at 4 °C, the upper supernatant and lipid layer were completely removed. The cyst-enriched fraction was collected, washed with 1× PBS, and centrifuged at 1580 g for 10 min. Finally, the pellet was resuspended in PBS for microscopic cyst quantification. All experiments were repeated three times.

### Purification Using Percoll Density Gradient Centrifugation

A three-layered discontinuous Percoll gradient (90%, 40%, and 20%, 2 mL per layer) was prepared. Brain homogenate was slowly layered on the surface of the 20% Percoll layer and centrifuged at 1,280 g for 25 min at 4 °C. The target fraction above the red blood cell layer was harvested, washed with 1× PBS, and centrifuged at 1,580 g for 10 min. The obtained pellet was resuspended in PBS for subsequent cyst counting.

### Purification Using CsCl Density Gradient Centrifugation

A four-step discontinuous cesium chloride (CsCl) density gradient with densities of 1.615, 1.308, 1.135, and 1.065 g/mL (2 mL per layer) was prepared in advance. A total of 1 mL of brain homogenate was layered onto the gradient surface and centrifuged at 2,800 × g for 45 min at 4 ℃. The upper supernatant and floating lipid layer were discarded. The precipitated pellet was washed with 1× PBS, centrifuged at 1,580 g for 10 min, and resuspended in PBS for cyst enumeration.

### In Vivo Infectivity Assay of Purified Cysts

Purified brain cysts were diluted to a concentration of 10 cysts per 200 μL of PBS. Mice were orally inoculated with the cyst suspension (n = 5 mice per group). All mice were euthanized 30 days post-inoculation, and brain tissues were collected and microscopically examined for the presence of *T. gondii* cysts.

### FDA/PI Double Fluorescence Staining for Cyst Viability

For viability assessment, 500 μL of purified cyst suspension was mixed with 1 μL of 5 mg/mL FDA working solution and 10 μL of 1 mg/mL PI working solution. The mixture was incubated at 37 ℃ for 20 min. Stained samples were observed under a fluorescence microscope, and cyst viability was evaluated based on fluorescence signals from no fewer than three random microscopic fields.

### Trypan Blue Exclusion Assay for Bradyzoite Viability

Purified intact cysts were digested with trypsin to release intracellular bradyzoites. The isolated bradyzoites were stained with 0.4% trypan blue solution for 5–10 min at 4 °C, followed by PBS washing to remove residual dye. Cell viability was observed and counted under a light microscope, with the survival rate calculated from no fewer than three independent microscopic fields.

### In Vivo Infectivity Assay of Isolated Bradyzoites

Trypsin-released bradyzoites were adjusted to a final concentration of 500 parasites per 200 μL of PBS. Mice received intraperitoneal injection of the bradyzoite suspension (n = 5 mice per group). Thirty days after inoculation, mice were euthanized, and brain tissues were isolated and examined microscopically to observe cyst formation and evaluate bradyzoite infectivity.

### Statistical analysis

Statistical analyses were performed using GraphPad Prism version 8.0.2. One-way analysis of variance (ANOVA) was applied to evaluate the statistical significance of differences. All data are expressed as mean ± standard deviation (SD) of at least three independent biological replicates. Statistical significance was defined as follows: ns, not significant (*P* > 0.05); \**P* < 0.05; \*\**P* < 0.01; \*\*\**P* < 0.001.

## Declarations

### Ethics approval and consent to participate

All animal procedures were approved by the Animal Ethics Committee of Beijing University of Agriculture (No. BUA2023118).

### Consent for publication

Not applicable

### Availability of data and materials

Not applicable

### Authors contributions

T.T.Y. carried out all experiments, analyzed the data and drafted the original manuscript. W.Y.M., X.F.L. and S.J.N. assisted with experimental procedures. H.R.L. and Q.M.L. offered overall supervision of the project. D.Q.Y. conceived the study design, conducted partial experiments and revised the manuscript. All authors have read and approved the final version of the manuscript.

## Acknowledgements

Not applicable

